# The Atypical Cyclin-Like Protein Spy1 Overrides p53-Mediated Tumour Suppression and Promotes Susceptibility to Breast Tumorigenesis

**DOI:** 10.1101/535310

**Authors:** Bre-Anne Fifield, Ingrid Qemo, Evie Kirou, Robert D. Cardiff, Lisa Ann Porter

## Abstract

**Background:** Breast cancer is the most common cancer to affect women and one of the leading causes of cancer related deaths. Proper regulation of cell cycle checkpoints plays a critical role in preventing the accumulation of deleterious mutations. Perturbations in the expression or activity of mediators of cell cycle progression or checkpoint activation represent important events that may increase susceptibility to the onset of carcinogenesis. The atypical cyclin-like protein Spy1 was isolated in a screen for novel genes that could bypass the DNA damage response. Clinical data demonstrates that protein levels of Spy1 are significantly elevated in ductal and lobular carcinoma of the breast. We hypothesized that elevated Spy1 would override protective cell cycle checkpoints and support the onset of mammary tumorigenesis.

**Methods:** We generated a transgenic mouse model driving expression of Spy1 in the mammary epithelium. Mammary development, growth characteristics and susceptibility to tumorigenesis was studied. *In vitro* studies were conducted to investigate the relationship between Spy1 and p53.

**Results:** We found that in the presence of wild-type p53, Spy1 protein is held ‘in check’ via protein degradation, representing a novel endogenous mechanism to ensure protected checkpoint control. Regulation of Spy1 by p53 is at the protein level and is mediated in part by Nedd4. Mutation or abrogation of p53 is sufficient to allow for accumulation of Spy1 levels resulting in mammary hyperplasia. Sustained elevation of Spy1 results in elevated proliferation of the mammary gland and susceptibility to tumorigenesis.

**Conclusions:** This mouse model demonstrates for the first time that degradation of the cyclin-like protein Spy1 is an essential component of p53-mediated tumour suppression. Targeting cyclin-like protein activity may therefore represent a mechanism of resensitizing cells to important cell cycle checkpoints in a therapeutic setting.

## Introduction

Breast cancer is the most prevalent form of cancer to affect women and represents the second leading cause of cancer related mortality among this population. Increased incidence of breast cancer in women can be attributed to the complex cellular changes the female mammary gland undergoes throughout life in response to hormonal cues. A delicate balance of cell cycle progression and inhibition is required at each of these periods of development to ensure maintenance of genomic stability; a crucial factor in the inhibition of tumourigenesis. Women with inherited mutations in genes that play fundamental roles in recognition of DNA damage and activation of DNA repair pathways have an elevated risk of breast cancer. Hence understanding how mammary epithelial cells monitor and respond to changes in genomic instability throughout development may reveal novel factors that predispose women to carcinogenesis.

The tumour suppressor p53 plays a critical role in DNA repair mechanisms, functioning to initiate arrest, repair and apoptotic programs [1–4]. Over 50% of human cancers contain a mutation in the *TP53* gene; individuals with Li-Fraumeni syndrome harbouring germline mutations in *TP53* are at an increased risk of developing cancer, including breast cancer, and mouse models with germline knockout of p53 develop normally however spontaneous tumours occur at an increased rate [5–10]. Thus, the inability of a cell to efficiently recognize and repair DNA damage plays a key role in the onset of tumourigenesis. Although p53 is widely mutated in human cancers and individuals with Li-Fraumeni syndrome have an elevated risk of breast cancer, this population comprises a small percentage of those with breast cancer, stressing the importance for cooperating genes in the initiation and/or progression of disease [11]. It is likely that these genes also play critical roles in normal cellular events that regulate proliferation, checkpoint activation and detection and repair of DNA damage, as aberrant expression of such genes would lead to genomic instability. Thus, it is of high importance to identify additional genes that may be implicated in breast cancer susceptibility.

An atypical cyclin-like protein Spy1 (also called Ringo, Speedy1; gene SPDYA) was initially discovered in a screen for genes that would override cell death following ultraviolet (UV) radiation in a rad1 deficient strain of *S. pombe*, suggesting a role for this protein in overriding critical checkpoint responses following DNA damage [12]. Several groups have demonstrated that Spy1 is capable of inhibiting apoptosis and promoting progression through both G1/S and G2/M phase of the cell cycle [13–16]. Spy1 function is currently attributed to the direct binding to the cyclin dependent kinases (Cdks), activating both Cdk1 and Cdk2 independent of threonine 161/160 phosphorylation status [14–19]. In the mammary gland, Spy1 protein levels are tightly regulated through development, being high during proliferative stages and downregulated at the onset of differentiation [20]. Interestingly, levels rise at the onset of involution, a period of development characterized by apoptosis and the triggering of regenerative processes [20]. When overexpressed in immortalized cells with a mutated p53 and transplanted in cleared fat pad assays, elevated levels of Spy1 protein leads to precocious development of the mammary gland, disrupts normal morphogenesis and accelerates mammary tumorigenesis [20]. Spy1 is elevated in human breast cancer [21, 22], as well as several other forms of cancer including brain, liver, and blood [23–25]. The ability of Spy1 to both enhance proliferation and override apoptosis and critical checkpoint responses provides further support for this finding. Spy1 may serve as an important mediator of the DNA damage response (DDR) in maintaining the proper balance of cellular proliferation; thus, deregulation of Spy1 may play a crucial role in the transition from precancerous to cancerous cell.

In this work we drive Spy1 overexpression in the mammary gland using the mouse mammary tumour virus (MMTV) promoter (MMTV-Spy1). We find that while glands are significantly more proliferative, there is no gross overall defect or pathology to the gland. Importantly, when hit with chemical carcinogens MMTV-Spy1 mice accumulate more DNA damage and have elevated susceptibility to mammary tumour formation. We noted that in this model endogenous wild-type-p53 was capable of keeping levels of Spy1 protein in check. We proceed to demonstrate a novel negative feedback loop with p53. This work demonstrates that tight regulation over the levels of cyclin-like proteins is a critical component of mammary tumour suppression and loss of control promotes hyperplastic growth and tumour initiation in the breast.

## Materials and Methods

### Construction of Transgene

The MMTV-Spy1 transgene was prepared as follows. Site directed mutagenesis was utilized to create an additional EcoRI site in Flag-Spy1A-pLXSN [26] to allow for subsequent removal of the Spy1 coding sequence using EcoRI digestion. The MMTV-SV40-TRPS-1 vector (kind gift from Dr Gabriel E DiMattia) was digested with EcoRI to remove the TRPS-1 coding sequence to allow for subsequent ligation of the Spy1 coding sequence into the MMTV-SV40 backbone.

### Generation and Maintenance of MMTV-Spy1 Transgenic Mice

MMTV-Spy1 (B6CBAF1/J-Tg(MMTV-Spy1)1Lport319, B6CBAF1/J-Tg(MMTV-Spy1)1Lport410, and B6CBAF1/J-Tg(MMTV-Spy1)1Lport413) mice were generated as follows: the MMTV-Spy1 vector was digested with XhoI and SpeI to isolate the MMTV-Spy1 transgene fragment and remove the remainder of the vector backbone. The transgene was sent to the London Regional Transgenic and Gene Targeting Facility for pronuclear injections in B6CBAF1/J hybrid embryos. Identification of founders and subsequent identification of positive pups was performed by PCR analysis. PCR was performed by adding 50 ng of genomic tail DNA to a 25µL reaction (1x PCR buffer, 2mM MgSO4, 0.2mM dNTP, 0.04U/µL BioBasic Taq Polymerase, 0.4µM forward primer [5’CCCAAGGCTTAAGTAAGTTTTTGG 3’], 0.4µM reverse primer [5’ GGGCATAAGCACAGATAAAACACT 3’], 1% DMSO) (NCI Mouse Repository). Cycling conditions were as follows: 94°C for 3 minutes, 40 cycles of 94°C for 1 minute, 55°C for 2 minutes, and 72°C for 1 minute, and a final extension of 72°C for 3 minutes. Mice were maintained hemizygously following the Canadian Council on Animal Care Guidelines under animal utilization protocol 14-22 approved by the University of Windsor.

### Primary Cell Harvest and Culture

Mammary tissue of the inguinal mammary gland was dissected and primary mammary epithelial cells were isolated as described [27]. Cells were also seeded on attachment plates in media containing 5% fetal bovine serum, 5 ng/mL EGF, 5 µg/mL insulin, 50 µg/mL gentamycin, 1% penicillin/streptomycin (P/S) in DMEM-F12 for BrdU incorporation assays conducted 1 week after isolation of the cells.

### Mammary Fat Pad Transplantation

The p53 knockout mouse, B6.129S2-Trp53tm1Tyj/J, was purchased from Jackson Laboratory (002101) [10]. Mammary epithelial cells were isolated from 8 week old mice and transplanted into the cleared glands of 3 week old B6CBAF1/J females. Successful clearing was monitored via the addition of a cleared gland with no injected cells.

### Cell Culture

Human embryonic kidney cells, HEK-293 (CRL-1573; ATCC), MDA-MB-231 (HTB-26; ATCC) and MCF7 (HTB-22; ATCC) were cultured in Dulbecco’s Modified Eagle’s Medium (DMEM; D5796; Sigma Aldrich) supplemented with 10% fetal bovine serum (FBS; F1051; Sigma Aldrich) and 10% calf serum (C8056; Sigma Aldrich) respectively, and 1% P/S. Mouse mammary epithelial cells, HC11 (provided by Dr. C. Shemanko) were maintained in RPMI supplemented with 10% newborn calf serum, 5 µg/mL insulin, 10 ng/mL EGF and 1% penicillin/streptomycin. All cell lines were maintained at 5% CO_2_ at 37°C. A BioRad TC10 Automated Cell Counter was used to assess cell viability via trypan blue exclusion. MG132 (Sigma Aldrich) was used at a concentration of 10 µM, and was added 12-16 hours post transfection. Cell lines purchased from ATCC were authenticated via ATCC. Cells were subject to routine mycoplasma testing. All cell lines were used within 3 passages of thawing.

### Plasmids

The pEIZ plasmid was a kind gift from Dr B. Welm, and the pEIZ-Flag-Spy1 vector was generated as previously described [24]. pCS3 and Myc-Spy1-pCS3 plasmids were generated as previously described [14], the Myc-Spy1-TST pCS3 plasmid was generated as previously described [28], and the p53-GFP backbone was purchased from Addgene (11770) (p53-GFP was a gift from Geoff Wahl (Addgene plasmid #11770)), (12091) (GFP-p53 was a gift from Tyler Jacks (Addgene plasmid #12091))[29]. The Nedd4DN vector was a kind gift from Dr. Dale S. Haines (Temple University School of Medicine). CMV10-3xFlag Skp2 delta-F was a gift from Sung Hee Baek (Addgene plasmid # 81116) [30].

### DMBA Treatments

Mice were given 1 mg of DMBA (Sigma Aldrich) in 100µL of a sesame:corn oil mixture (4:1 ratio) via oral gavage once per week. Treatment began when mice reached 8 weeks of age and were continued for 6 consecutive weeks. Mice were monitored on a weekly basis for the presence of tumours via palpitations. Mice were humanely sacrificed when tumours were noted, and all mice were sacrificed by 8 months of age regardless of tumour formation. Tissues were collected from sacrificed mice and flash frozen for immunoblotting and qRT-PCR analysis, or fixed in formalin for immunohistochemistry. DMBA was dissolved in DMSO for all *in vitro* experiments and used at a final concentration of 1.5 µg/mL.

### Histology and Immunostaining

Tissue was collected and fixed in 10% neutral buffered formalin. Immunohistochemistry was performed as described [31]. All primary antibodies were diluted in 3% BSA-0.1% Tween-20 in 1x PBS with the exception of mouse antibodies, which were diluted with Mouse on Mouse (MOM) blocker (Biocare Medical). Primary antibodies used were as follows: Spy1 (1:200; PA5-29417; Thermo Fisher Scientific), BrdU (1:200; 555627; BD Bioscience), γH2AX (1:200; 05-636; Millipore) Nedd4 (1:200; MBS9204431; MyBioSource), PCNA (1:500; sc-9857; Santa Cruz), and cleaved-caspase 3 (1:250; 9661; Cell Signaling). Secondary antibodies were used at a concentration of 1:750 and were as follows: Biotinylated anti-mouse, biotinylated anti-goat and biotinylated anti-rabbit (Vector Laboratories). Slides were imaged using the LEICA DMI6000 inverted microscope with LAS 3.6 software.

### Whole Mount Analysis

Briefly, the inguinal mammary gland was spread onto a positively charged slide (Fisherbrand 12-550-15) and left in Clarke’s Fluid (75% ethyl alcohol, 25% acetic acid) overnight. The following day, glands were placed in 70% ethyl alcohol for 30 minutes before being stained in carmine alum (0.2% carmine, 0.5% potassium aluminum sulphate) overnight. Glands were destained for 4 to 6 hours with destaining solution (1% HCl, 70% ethyl alcohol) and subsequently dehydrated in ascending concentrations of alcohol (15 minutes each 70, 95, 100% ethyl alcohol) before being cleared in xylene overnight. Slides were mounted with Permount toluene solution (Fisher Scientific) before imaging on a Leica MZFLIII dissecting microscope (University of Windsor). Images were captured using Northern Eclipse software.

### Transfection and Infection

MDA-MB-231 and MCF7 mammary cell lines were transiently transfected in serum and antibiotic free media using 25 µg of polyethylenimine (PEI) and 12 ug of plasmid DNA, incubated at room temperature for 10 minutes in base media before being added to the plate. For transfection of HC11 cells, media was changed to serum and antibiotic free media 4 hours prior to transfection. After 4 hours, 28 ug of PEI and 12 ug of plasmid DNA were incubated at room temperature for 10 minutes in base media before being added to the plate. Transfection of HEK-293 cells was performed in full growth media with 25 ug of PEI and 10 ug of plasmid DNA. For all cell lines transfection reagent was left for 16-18 hours.

Transfection of primary mouse cell lines with sip53 (Santa Cruz) and siRNA control (Santa Cruz) was performed using siRNA Transfection Reagent (Santa Cruz) as per manufacturer’s instructions.

### UV Irradiation

Media was removed from exponentially growing cells and cells were washed once with 1X PBS and subjected to 254 nm of UV radiation using a GS Gene Linker (Bio Rad). Immediately following irradiation, fresh medium was added to the cells.

### Quantitative Real Time PCR Analysis

RNA was isolated using Qiagen RNeasy Plus Mini Kit as per manufacturer’s instructions. cDNA was synthesized using Superscript II (Invitrogen) as per manufacturer’s instructions. SYBR Green detection (Applied Biosystems) was used for real time PCR and was performed and analyzed using Viia7 Real Time PCR System (Life Technologies) and software.

### Protein Isolation and Immunoblotting

Tissue lysis buffer (50mM Tris-HCl pH 7.5, 1% NP-40, 0.25% Na-deoxycholate, 1mM EGTA, 0.2% SDS, 150mM NaCl) with protease inhibitors (leupeptin 2 μg/mL, aprotinin 5 μg/mL, PMSF 100 μg/mL) was added to flash frozen tissue. Tissue and lysis buffer were homogenized on ice using a Fisher Scientific Sonic Dismembrator 50. Samples were centrifuged at 13000rpm for 20 minutes at 4°C. Supernatant was collected and centrifuged again at 13000rpm for 20 minutes at 4°C. Supernatant was collected and stored at −20°C until future use. Cells were lysed with TNE buffer (50 mM Tris, 150mM NaCl, 5mM EDTA) with protease inhibitors (leupeptin 2 μg/mL, aprotinin 5 μg/mL, PMSF 100 μg/mL). Cells were lysed for 10 minutes on ice, centrifuged at 4°C at 10,000rpm for 10 minutes, and supernatant was collected and stored at −20°C until further use.

Protein concentrations were assessed using the Bradford assay as per manufacturer’s instructions. Equal amounts of protein were analyzed and separated using SDS-PAGE and transferred to PVDF membranes. Membranes were blocked for 1 hour at room temperature in 1% BSA and incubated in primary antibody overnight at 4°C. Primary antibodies were used at a concentration of 1:1000 and were as follows: Actin (MAB1501; Millipore), p53 (ab131442; Abcam), Spy1 (ab153965; Abcam), c-Myc (C3956; Sigma Aldrich), Flag (F1804; Sigma Aldrich), Nedd4 (MBS9204431; MyBioSource). Secondary antibody mouse or rabbit IgG (Sigma Aldrich) at a concentration of 1:10,000 was used for 1 hour at room temperature. Visualization was conducted using chemiluminescent peroxidase substrate (Pierce) as per manufacturer’s instructions. Images were captured on Alpha Innotech HD 2 using AlphaEase FC software.

### BrdU Incorporation Assay

15,000 cells per well were seeded in a 96 well plate. BrdU (BD Pharmingen) was added 24 hours later to a final concentration of 10 µM and cells were incubated in media containing BrdU for 24 hours at 37°C, 5% CO_2_. Media containing BrdU was removed and cells were washed three times with 1x PBS. Cells were fixed in 4% PFA for 15 minutes, washed twice with 1xPBS, incubated for 20 minutes at 37°C in 2M HCl and subsequently washed once with 1x PBS. Cells were incubated for 45 minutes with Anti-BrdU (BD Biosciences) in 0.2% Tween in 1x PBS. Cells were washed with 1x PBS and incubated with anti-mouse IgG and hoescht at a 1:1000 dilution in 1x PBS for 1 hour at room temperature. Cells were washed one time with 1x PBS, once with distilled water and stored at 4°C in 50% glycerol until imaged using the LEICA DMI6000 inverted microscope.

### Flow Cytometry

Mammary primary epithelial cells were isolated from inguinal glands as described (27). Cells were stained using CD24 (APC; BD 562349) and CD45 (PeCy7; BD 552848), and FACS was performed using a BD LSR Fortessa X-20 (Becton Dickinson).

### Statistical Analysis

For tumour studies, a Mann-Whitney test was performed for statistical analysis. For all other data, a Student’s T-Test was performed. Unequal variance was assumed for experiments involving mouse tissue samples and primary mammary epithelial cells. Cell line data analysis assumed equal variance. All experiments, both *in vitro* and *in vivo*, included at least 3 biological replicates and results are representative of at least 3 experimental replicates. No randomization or blinding occurred for animal studies. Significance was scored as *p<0.05, **p<0.01, ***p<0.001.

See supplemental information for more materials and methods.

## Results

### Generation of MMTV-Spy1 transgenic mice

The flag-Spy1 coding sequence was cloned into the MMTV-SV40 plasmid (Figure 1A) and injected into B6CBAF1/J pronuclei. PCR analysis identified three founders, with 5 to 15 copies of the transgene (data not shown), all of which successfully transmitted the transgene to their progeny (Figure S1A). Analysis of both mRNA and protein levels from 6-week-old mice revealed that mammary glands from MMTV-Spy1 mice contained significantly higher levels of Spy1 as compared to control littermates (Figure S1B). Western blot analysis of other tissues in the MMTV-Spy1 mice did not demonstrate significant elevation of Spy1 (Figure S1C).

**Figure 1:**
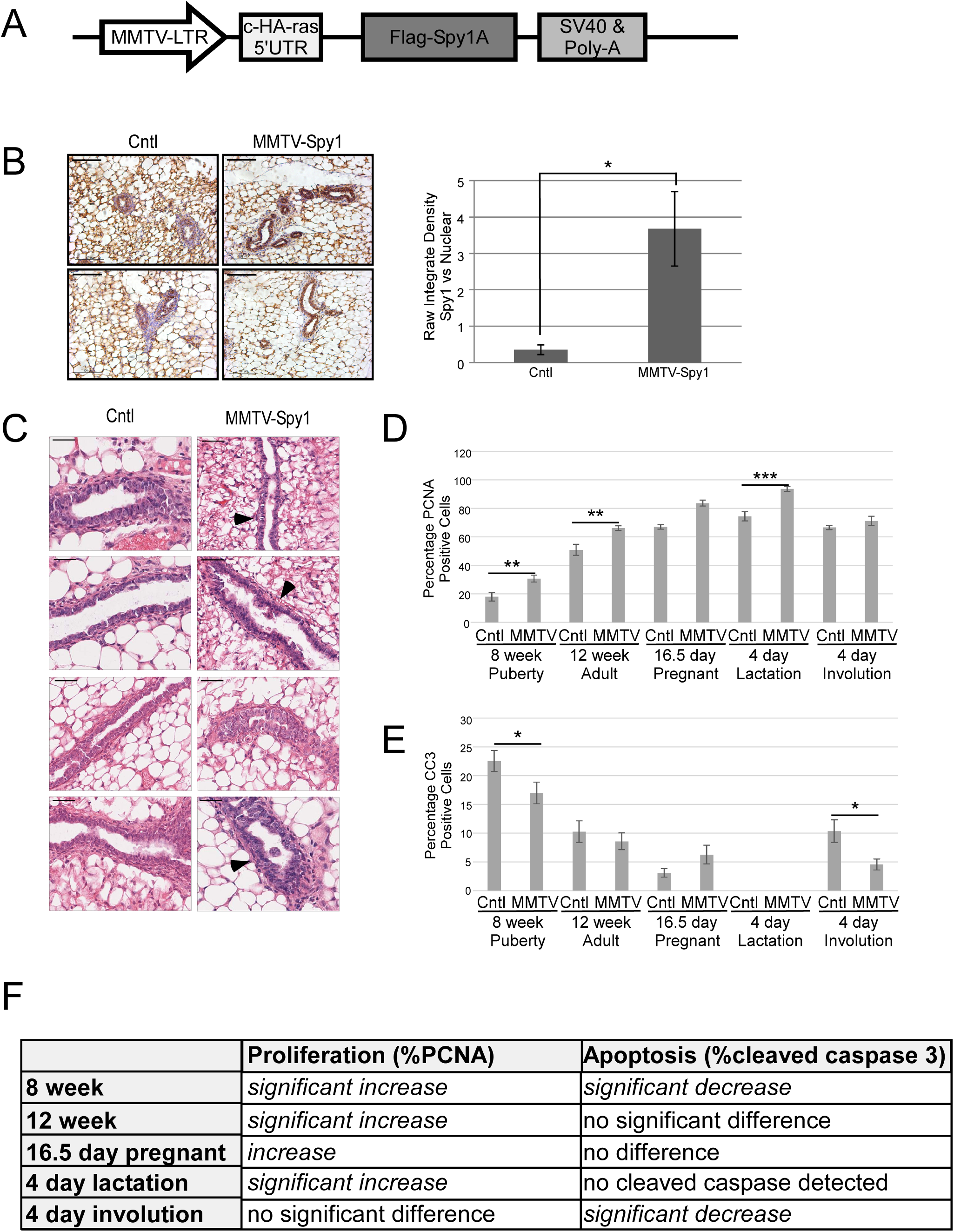
Characterization of MMTV-Spy1 mice. A) Schematic representation of the MMTV-Spy1 transgenic vector used in pronuclear injections for the generation of the MMTV-Spy1 mouse. B) Spy1 expression in 8-week-old MMTV-Spy1 and control littermate (Cntl) inguinal mammary glands, where blue stain is haematoxylin and brown stain represents Spy1 expression. Representative images in left panels with quantification of Spy1 levels using ImageJ software analysis shown in right panel. Scale bar= 100 µm. C) Representative H&E stain of inguinal mammary glands from 6-week-old MMTV-Spy1 mice and control littermates (Cntl). Scale bar = 50 µm. D) PCNA expression in MMTV-Spy1 and littermate controls via immunohistochemical analysis. Quantification of percentage of PCNA positive mammary epithelial cells over 5 fields of view per sample (8 week Cntl n=3, MMTV-Spy1 n=4; 12 week Cntl n=3, MMT-Spy1 n=3; 16.5 day pregnant Cntl n=1, MMTV-Spy1 n=2; 4 day lactation Cntl n=3, MMTV-Spy1 n=2; 4 day involution Cntl n=2, MMTV-Spy1 n=2). E) Cleaved caspase 3 (CC3) expression in MMTV-Spy1 and littermate controls via immunohistochemical analysis. Quantification of percentage of CC3 positive mammary epithelial cells over 5 fields of view per sample (8 week Cntl n=3, MMTV-Spy1 n=4; 12 week Cntl n=3, MMT-Spy1 n=3; 16.5 day pregnant Cntl n=1, MMTV-Spy1 n=2; 4 day lactation Cntl n=3, MMTV-Spy1 n=2; 4 day involution Cntl n=2, MMTV-Spy1 n=2). F) Summary of proliferation and apoptosis data for developmental time course. Error bars reflect standard error (SE), Student’s T-test *p<0.05, **p<0.01, ***p<0.001. See also Figures S1 and S2.

Previous data demonstrated that increased levels of Spy1 in immortalized ‘normal’ mouse mammary cells (HC11 cells) transplanted into cleared fat pads can disrupt morphology of the mammary gland and promote accelerated development *in vivo* [20]. Histopathological analysis of MMTV-Spy1 glands during puberty revealed modest phenotypic changes in the gland including a thickening of the ductal walls and some abnormal, proliferative characteristics (Figures 1C black arrowheads). Additionally, Spy1 appeared to be expressed primarily in luminal cells and showed varying expression in myoepithelial cells (Figures 1B, S1D). Flow cytometry was used to delineate between basal and luminal populations of cells as described [32] and while there does appear to be increases in epithelial content, no significant difference was observed (Figure S1E). Gross morphology of the gland was not altered in whole mount analysis or histological analysis at any developmental time point analysed (Figures S2A,B,C). All MMTV-Spy1 female mice successfully nursed their litters, even following multiple rounds of pregnancy and there were no tumours noted when mice were aged for 2 years.

Spy1 increases cell proliferation in a variety of cell types when exogenously expressed [14, 22]. To determine if MMTV-Spy1 mammary glands exhibited increased rates of proliferation, immunohistochemical analysis was performed to examine the expression of PCNA throughout a developmental time course. MMTV-Spy1 mice had significantly more PCNA positive cells than their littermate controls indicating increased proliferation at all points examined except for day 4 of involution (Figure 1D,F,S3). To determine if there was a bone fide increase in proliferation with no subsequent increase in apoptosis to counterbalance enhanced proliferation, glands were analyzed for expression of cleaved-caspase 3. No differences in cleaved-caspase 3 were detected at 12 weeks, day 16.5 pregnancy or during lactation; however, a significant reduction in apoptosis was seen at 8 weeks and day 4 of involution (Figure 1E,F,S3). This suggests that Spy1 is capable of not only enhancing proliferation but also overriding apoptosis in an *in vivo* setting. To further validate this finding, primary mammary epithelial cells were isolated from the inguinal mammary glands of control and MMTV-Spy1 mice and treated with BrdU. Cells from MMTV-Spy1 inguinal mammary glands were found to have a significantly higher percentage of BrdU positive cells (Figure S2D). Hence, MMTV-Spy1 mice display modest phenotypic and no gross morphological changes in the mammary gland despite having enhanced proliferation and decreased apoptosis.

### Spy1 increases mammary tumour susceptibility

Although MMTV-Spy1 mammary glands exhibit significant changes in proliferative capacity, they develop normally and do not present with spontaneous tumours. Increased protein levels of Spy1 have been implicated in several human cancers including that of the breast, ovary, liver and brain [20, 22–24]. To assess whether or not elevated levels of Spy1 may affect tumour susceptibility, MMTV-Spy1 mice and control littermates were treated with the mammary carcinogen 7,12-dimethlybenz(a)anthracene (DMBA) once per week for 6 consecutive weeks during puberty (Figure 2A). DMBA induces DNA damage through the formation of DNA adducts and is commonly used in rodent models to study the onset and timing of mammary tumour formation [33–35]. Mice were monitored on a weekly basis for tumour formation. The timing of tumour initiation was not altered (Figure 2B) however, 95% of MMTV-Spy1 mice developed tumours as compared to only 45% of control mice (Figure 2C). Of the tumours developed, 80% of MMTV-Spy1 mice presented with mammary tumours both benign and malignant, as compared to only 30% of littermate controls. Interestingly, ovarian tumours occurred in MMTV-Spy1 mice, but there was no incidence of ovarian tumours in littermate controls. Tumour tissue was sent for pathological analysis, and MMTV-Spy1 mice had significantly more malignant mammary tumours over littermate controls (Figures 2D-E).

**Figure 2:**
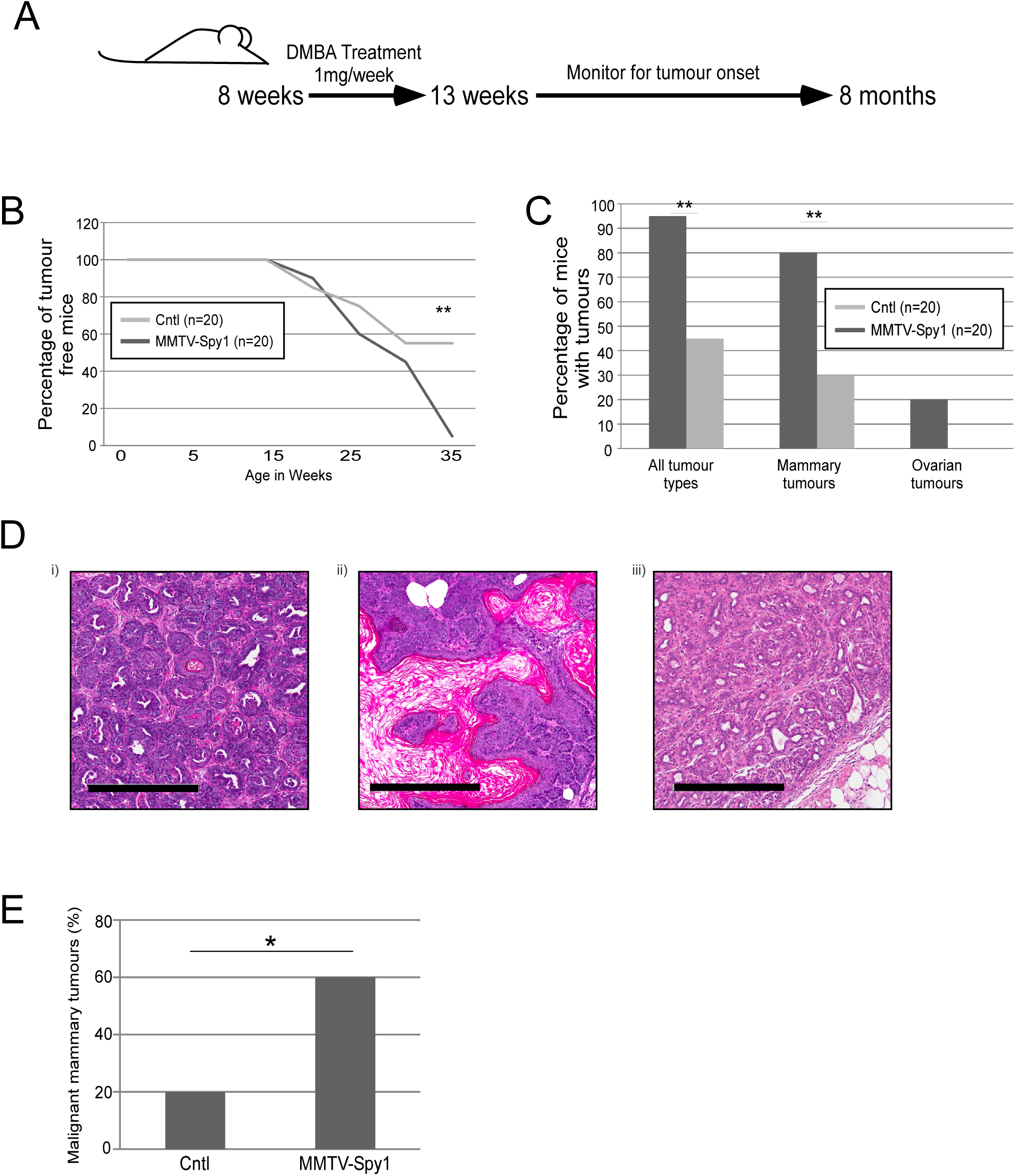
Spy1 overexpression leads to increased mammary tumour susceptibility. A) Schematic of DMBA treatment. B) Graphical representation of timing of tumour onset (n=20). C) Graphical representation of percentage of mice with tumours (n=20). D) Representative images of tumour pathology from DMBA induced mammary tumours i) and ii) adenosquamous carcinoma, iii) adenomyoepithelioma. Scale bar= 300 µm. E) Graphical representation of the number of mice with malignant mammary tumours (n=20). Mann-Whitney*p<0.05, **p<0.001

### p53 can regulate protein levels of Spy1

Previous mammary fat pad transplantation of Spy1 overexpressing HC11 cells leads to increased tumour formation *in vivo* [20]. HC11 is an immortalized cell line with mutated p53 that renders p53 non-functional [36–38]. Spy1 is capable of preventing checkpoint activation [15] and since p53 plays a critical role in mediating proper checkpoint activation, it is plausible then that the lack of spontaneous tumours in the MMTV-Spy1 mice may be attributed to the presence of wild-type p53. To test this theory, primary mammary epithelial cells were extracted from an MMTV-Spy1 mouse and p53 was knocked down using siRNA (Figure 3A). Interestingly, with only a modest decrease in p53 protein levels (Figure 3A; middle panel) there was a very significant increase in Spy1 protein levels (Figure 3A; left panel). Given that tumour formation was seen in a cell line with non-functional p53, and Spy1 can prevent checkpoint activation [13, 15, 16, 20], it is plausible then that wild-type p53 may work to downregulate Spy1 to allow for p53 mediated cell cycle arrest, and elevated Spy1 with loss of p53 function would allow for enhanced genomic instability. To test the ability of wild-type p53 to regulate levels of Spy1 protein, mammary cells with mutated p53 (HC11 and MDA-MB-231 cells) were transfected with pEIZ, pEIZ-Spy1, p53 or pEIZ-Spy1 and p53 and lysates collected at 24 hours for Western blot analysis. Levels of Spy1 protein were significantly decreased in the presence of wild-type p53 (Figure 3B). To determine if p53 also affected Spy1 mRNA, MDA-MB-231 cells were transfected with pEIZ, pEIZ-Spy1, p53 or pEIZ-Spy1 and p53 and levels of mRNA were assessed via qRT-PCR. There was no effect on levels of Spy1 mRNA in the presence of elevated p53 indicating that p53 likely regulates Spy1 expression at the level of protein expression (Figure S4A). Previous data has demonstrated that Spy1 is targeted for proteasome-dependent degradation via either the E3 ubiquitin ligase Nedd4 [28] or the Skp2 ubiquitin ligase [39], which pathway is active may depend on phase of the cell cycle. To determine if the downregulation of Spy1 by p53 is proteasome dependent, Spy1 and p53 were expressed in the presence of the proteasome inhibitor MG132. Inhibition of the proteasome in the presence of p53 abrogated the downregulation of Spy1 protein, supporting that p53 regulates protein levels of Spy1 via a proteasome dependent mechanism (Figure 3C). To determine whether Nedd4 or Skp2 were responsible for p53-mediated degradation of Spy1, Spy1 and p53 were overexpressed along with dominant negative forms of both Nedd4 and Skp2. Levels of Spy1 were significantly decreased in the presence of p53 and the dominant negative Skp2; however, loss of Nedd4 activity significantly reduced the ability of p53 to decrease levels of Spy1 (Figure 3D). To determine if p53 is capable of mediating levels of Nedd4, p53 was overexpressed and protein and RNA levels of Nedd4 were examined. No significant differences were seen at either the levels of protein or RNA (Figure S4B,C). Previous data has also demonstrated that post-translational modification of Spy1 at residues Thr15, Ser22, Thr33 targets Spy1 for degradation by Nedd4 [28]. Wild-type Spy1 and a mutant non-degradable by Nedd4 (Spy1-TST) were both overexpressed in the presence of p53. Levels of wild-type Spy1 are significantly decreased in the presence of p53; however, p53 is unable to downregulate Spy1-TST indicating that post-translational modifications of Spy1 play an important role in p53 mediated degradation of Spy1 (Figure 3E). This data supports that Spy1 levels are tightly controlled by p53 and this response is dependent on the E3 ligase Nedd4.

**Figure 3:**
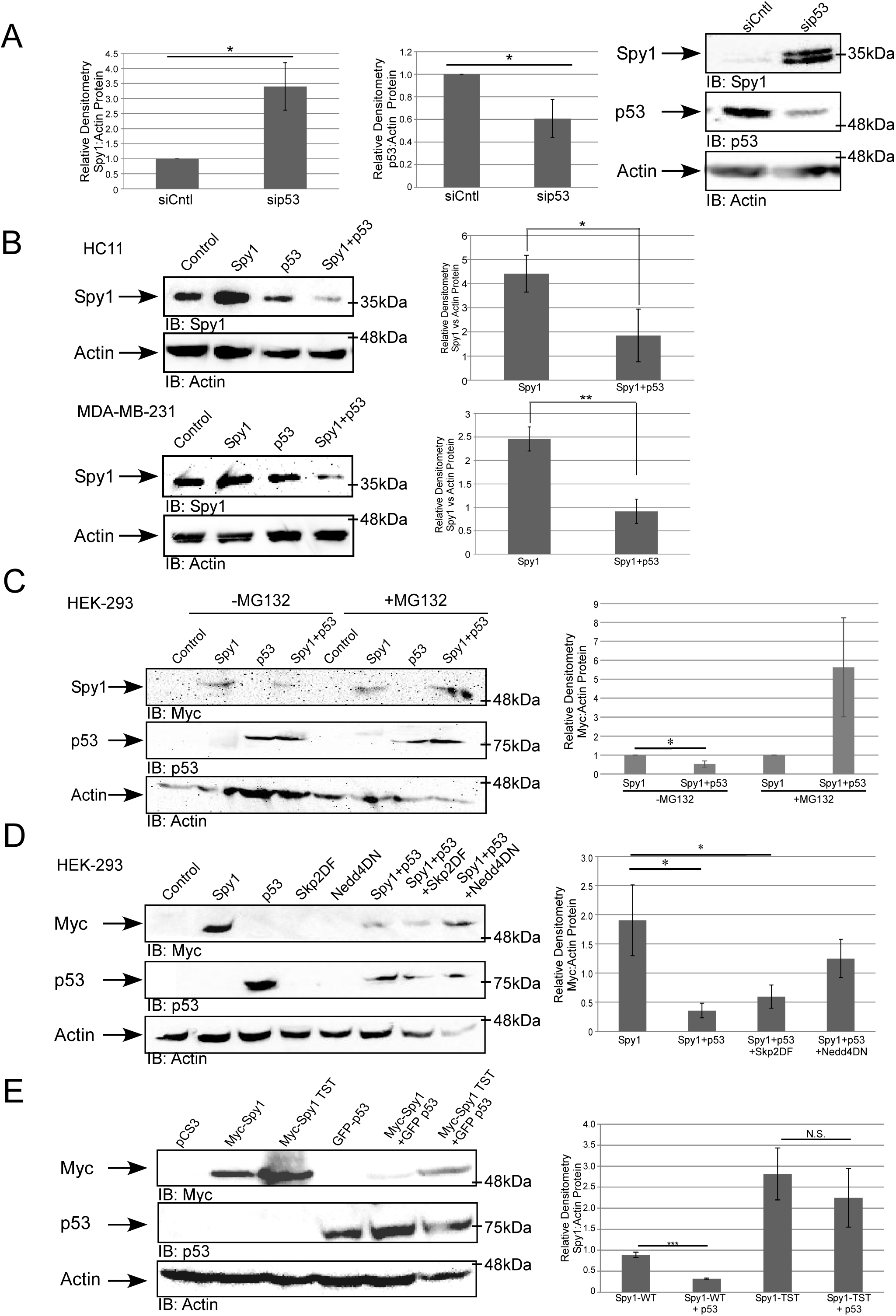
p53 regulates Spy1 protein levels through the ubiquitin ligase Nedd4. A) Western blot analysis of Spy1 (left panel) and p53 (middle panel) protein levels in MMTV-Spy1 primary mammary epithelial cells corrected for Actin. Data is represented as a fold change as compared to control siRNA (siCntl). Representative blot is shown in right panel. B) Levels of Spy1 protein were assessed via Western blot analysis 24 hours after transfection in HC11 (n=6) and MDA-MB 231 (n=5) cells transfected with pEIZ, pEIZ-Spy1, p53 or both pEIZ-Spy1 and p53. Left panels depict representative blots and right panels depict densitometry analysis of Spy1 levels corrected for Actin. C) Levels of Spy1 protein were assessed via Western blot analysis in presence and absence of MG132. Left panel depicts representative blot and right panel depicts densitometry analysis of Spy1 protein levels corrected for Actin. Data is shown as fold change to cells transfected only with the Spy1 vector (n=3). D) Levels of Spy1 protein were assessed in HEK-293 cells after transfections with control vector pCS3 and Myc-Spy1-pCS3, p53, Skp2ΔF, and Nedd4DN in various combinations. Cells were collected 24 hours after transfection and subjected to Western blot analysis. Densitometry analysis was performed for total Spy1 protein levels and corrected for total Actin levels (n=3). E) Levels of Spy1 and Spy1-TST protein were assessed in HEK-293 cells after transfection with control vector pCS3, myc-Spy1-pCS3, myc-Spy1-TST-pCS3 and p53. Cells were collected 24 hours after transfection and subjected to Western blot analysis. Densitometry analysis was performed for total Spy1 protein levels and corrected for total Actin levels (n=3). Errors bars represent SE; Student’s T-test. *p<0.05, **p<0.01, ***p<0.001, not significant (N.S.). See also Figure S3.

### Spy1 downregulation is a necessary component of the DDR

Spy1 can override the function of downstream effectors of p53 [13, 15], hence we hypothesize that negative regulation of Spy1 by wild-type p53 may be essential to ensure a healthy DDR response. To test this, cell proliferation was measured in HC11, MCF7 and MDA-MB-231 cells following Spy1, p53 or Spy1 and p53 overexpression in the presence or absence of DNA damage stimuli (Figures 4A-B). Spy1 was capable of overriding the effects of constitutive expression of p53 both in the presence and absence of damage in both DMBA (Figure 4A) and UV damage (Figure 4B). It is notable that this effect was independent of endogenous p53 status. To further examine the functional relationship between Spy1 and p53 in primary mammary epithelial cells, p53 levels were manipulated with siRNA in cells extracted from the MMTV-Spy1 mice or littermate controls (Figure 4C; left panel). Cell proliferation was measured in the presence and absence of UV damage (Figure 4C; right panel). These data demonstrate that endogenous levels of wild-type p53 keep a check on primary mammary populations in both the presence and absence of damage and that loss of p53 resulted in a robust increase in Spy1-mediated effects on proliferation.

**Figure 4:**
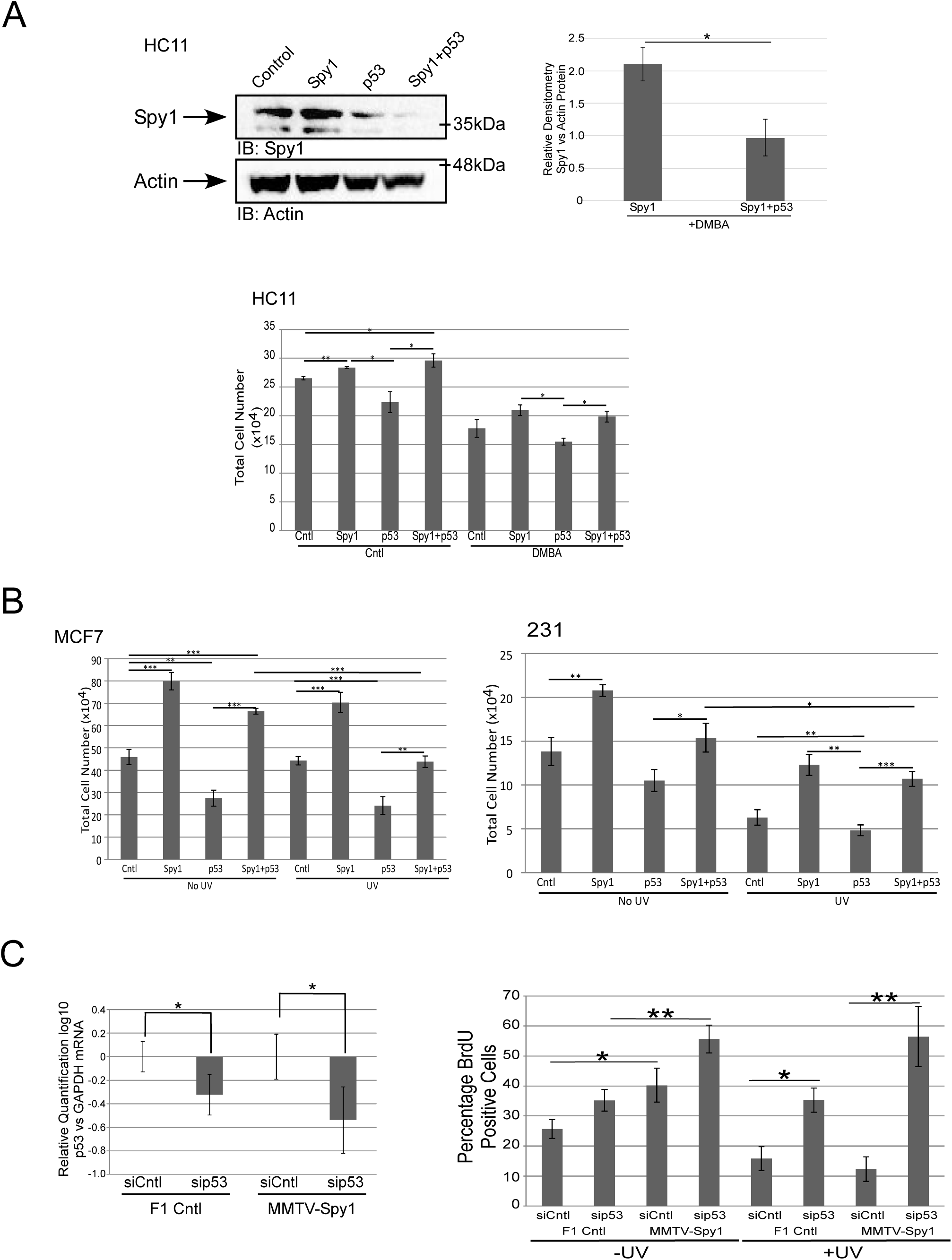
Spy1 can enhance proliferation in the presence of p53. A) HC11 cells were transfected with vector control, pEIZ-Spy1, p53 or pEIZ-Spy1 and p53 in the presence or absence of 1.5µg/mL of DMBA. Levels of Spy1 are depicted (upper panels). Growth of cells following transfection was assessed via trypan blue analysis (lower panels) (n=3). B) MCF7 (left panel) and MDA-MB 231(right panel) were transfected with vector control, pEIZ-Spy1, p53 or pEIZ-Spy1 and p53 in the presence or absence of 50J/m^2^ UV damage. Growth of cells following transfection was assessed via trypan blue analysis (n=3). C) qRT-PCR analysis of p53 levels in littermate control (F1 Cntl) and MMTV-Spy1 primary mammary epithelial cells corrected for total GAPDH. (left panel). Quantification of BrdU positive cells with and without UV irradiation with (siCntl) and without p53 (sip53) (right panel). F1 Cntl n=5, MMTV-Spy1 n=5. Error bars represent SE; Student’s T-test. *p<0.05, **p<0.01, ***p<0.001.

### Spy1 expression disrupts the DDR in the presence of DMBA

To validate the *in vitro* findings that Spy1 elevation can alter proper checkpoint activation, MMTV-Spy1 mice were treated with 1 mg DMBA, and inguinal mammary gland tissues were collected after 48 hours and analysed for alterations in known DDR proteins (Figure 5A). Spy1 was significantly overexpressed at the mRNA level in 8-week-old MMTV-Spy1 mice with and without DMBA (Figure S5A). Spy1 protein levels were also elevated in the MMTV-Spy1 mice over littermate controls both in the presence and absence of DMBA (Figure 5B; left panel). Importantly, Spy1 protein levels increased in control mice following treatment with DMBA in accordance with previous data demonstrating Spy1 is upregulated in response to damage [15]. Interestingly, p53 levels were significantly higher in the MMTV-Spy1 mice over littermate controls after DMBA treatment (Figure 5B compare left to right panels, Figure S5B). MMTV-Spy1 mice treated with DMBA were also found to have a significant increase in Nedd4 expression at the same time as p53 suggesting an upregulation in pathways responsible for Spy1 mediated degradation (Figure 5C).

**Figure 5:**
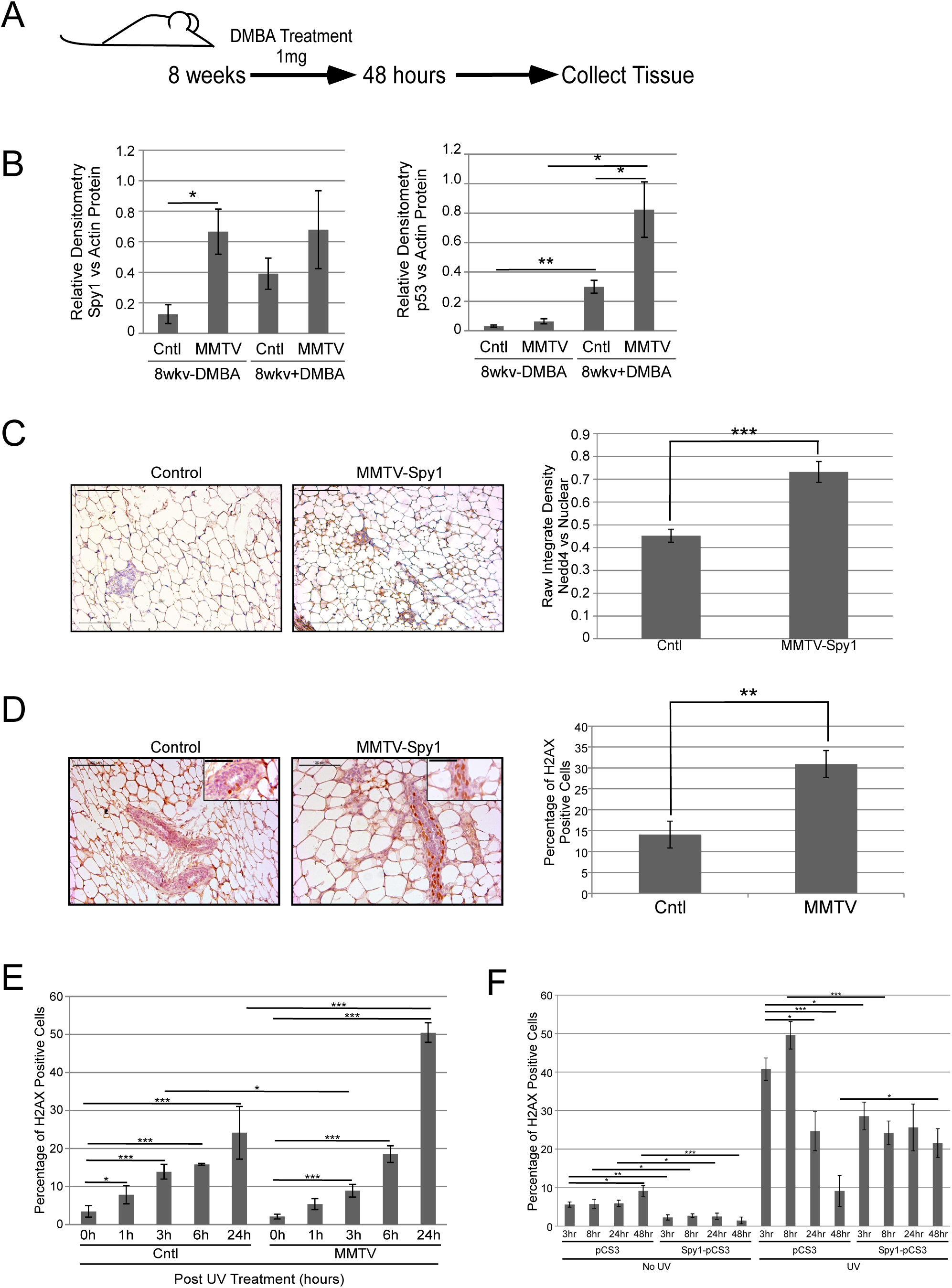
MMTV-Spy1 mice show alterations in DDR pathway when exposed to DMBA. A) Schematic of short term DMBA treatment and collection of samples. B) Western blot for Spy1 (left panel) and p53 (right panel) levels in 8-week-old control mice and DMBA treated mice 48 hours following DMBA exposure. Densitometry analysis is depicted with total Spy1 and p53 levels corrected for total levels of Actin. C) Immunohistochemical analysis for Nedd4 expression in inguinal mammary glands of 8-week-old MMTV-Spy1 mice and littermate controls was performed after exposure to DMBA. Representative images are shown in left panel. Levels of Nedd4 were quantified using ImageJ analysis (right panel). Scale bar=100 µm D) Representative images of immunohistochemical analysis of γH2AX in inguinal mammary glands of 8-week-old MMTV-Spy1 and littermate control (Cntl) mice after exposure to DMBA (left panel), where brown stain is γH2AX and blue stain is haematoxylin. Number of γH2AX positive cells were counted and quantified as percentage of γH2AX cells (right panel). Scale bars= 100 µm and 50 µm (inset image) E) Primary mammary epithelial cells from MMTV-Spy1 mice and control littermates were isolated and UV irradiated with 50J/m^2^. Cells were collected 0, 1, 3 6 and 24 hours post-UV and immunofluorescence was performed to assess formation of γH2AX foci following damage (n=3). F) HC11 cells were transfected with pCS3 and Myc-Spy1-pCS3, and UV irradiated with 50J/m^2^. Cells were analysed at various times following UV irradiation for the number of γH2AX positive cells via immunofluorescence. Errors bars represent SE; Student’s T-test. *p<0.05, **p<0.01, ***p<0.001.

### Elevated levels of Spy1 lead to accumulated DNA damage

The effects of Spy1 on the level of DNA damage following exposure to DMBA was investigated *in vivo*. MMTV-Spy1 mice at 8 weeks of age were again treated once with DMBA and samples were collected and analyzed 48 hours post treatment. MMTV-Spy1 mice had significantly more γH2AX positive cells as compared to littermate controls, indicating a lack of repair in response to DMBA (Figure 5D). To determine if this is ubiquitous for different forms of DNA damage, primary inguinal mammary gland cells from MMTV-Spy1 mice and control littermates were isolated and UV irradiated with 50 J/m^2^. Expression of γH2AX was monitored at a time course following damage. Cells from MMTV-Spy1 mice had significantly more γH2AX positive cells at 24 hours post UV as compared to control littermate cells (Figure 5E). Data from the MMTV-Spy1 mouse both *in vivo* and *in vitro* shows a significant increase in γH2AX following DNA damage, which is in opposition to previously published data, which shows a significant decrease in γH2AX with Spy1 overexpression [13, 16]. To determine if this is due to a difference in the time points studied, HC11 cells were transfected with pCS3 or Myc-Spy1-pCS3, UV irradiated and studied at a wide time course. At all times collected in non-irradiated cells, Spy1 overexpression led to a significant decrease in γH2AX as compared to control (Figure 5F). Following UV however, γH2AX was significantly lower in Spy1 cells at early time points and then significantly higher at 48 hours post UV. Previous work has examined the role of Spy1 in checkpoint activation following damage [13, 16]. Spy1 overexpression lead to decreased activation of both S phase and G2M checkpoints, as well as decreased activation of DDR signaling as assessed through Chk1 phosphorylation status [13, 16]. Spy1 also decreased rates of removal of damage following UV, indicating that elevated levels of Spy1 prevent cellular checkpoint activation and impair removal of damage [13]. This data supports that elevated levels of Spy1 may promote proliferation and a delayed or impaired recognition of DNA damage at early time points, however overriding checkpoints over time leads to an accumulation of DNA damage.

### In the absence of p53, Spy1 drives hyperplasia

To determine if loss of p53 cooperates with Spy1 to promote tumourigenesis, levels of p53 were assessed in DMBA treated MMTV-Spy1 mice and their control littermates at end point (Figure 2A) to determine if a decrease in p53 correlated with the development of tumours in DMBA treated MMTV-Spy mice. Levels of p53 were significantly lower in both MMTV-Spy1 DMBA induced mammary tumours as well as surrounding normal mammary tissue as compared to control (Figure 6A). Interestingly, there was no difference in p53 expression in control surrounding normal mammary tissue as compared to control DMBA mammary tumours, while MMTV-Spy1 DMBA mammary tumours had significantly lower p53 as compared to MMTV-Spy1 normal mammary tissue (Figure 6A).To determine if the loss of p53 is sufficient to drive spontaneous tumourigenesis with elevated levels of Spy1, MMTV-Spy1 mice were crossed with p53 null mice. Mammary fat pad transplantation was performed when mice were 8 weeks of age to transplant extracted primary mammary epithelial cells from the resulting crosses into the cleared fat pad of 3 week old wild type mice to eliminate the possibility of other tumours forming prior to the onset of mammary tumours. Mice were left to age for up to 2 years and monitored for formation of spontaneous mammary tumours. Whole mount analysis was performed on glands that did not develop tumours to assess for the formation of hyperplastic alveolar nodules (HANs) (Figure 6B, C). There was a significant increase in formation of HANs and tumours in fat pads of wild-type mice reconstituted with mammary epithelial cells from intercrossed MMTV-Spy1 p53-/-mice as compared to mice reconstituted with wild-type mammary epithelial cells. One MMTV-Spy1 p53+/-mouse developed a mammary tumour at 25 weeks post-transplant, while no p53+/-mice developed tumours even when left to 2 years of age. Two p53-/-and two MMTV-Spy1 p53-/-mice developed tumours and there was no difference in number of glands with HANs or tumours when comparing p53+/-to MMTV-Spy1 p53+/-. Complete loss of p53 with elevated levels of Spy1 lead to increased formation of HANs when comparing p53 loss alone with p53 loss combined with elevated Spy1 (Figure 6B). Numbers of both p53 -/-and MTMV-Spy1 p53-/-were lower than expected Mendelian ratios likely due to embryonic lethality. Elevated levels of Spy1 appear to enhance hyperplastic growth of mammary gland tissue when combined with loss of p53. This data supports the conclusion that wild-type p53 holds Spy1 levels in check to permit successful checkpoint regulation and preserve genomic integrity of the gland.

**Figure 6:**
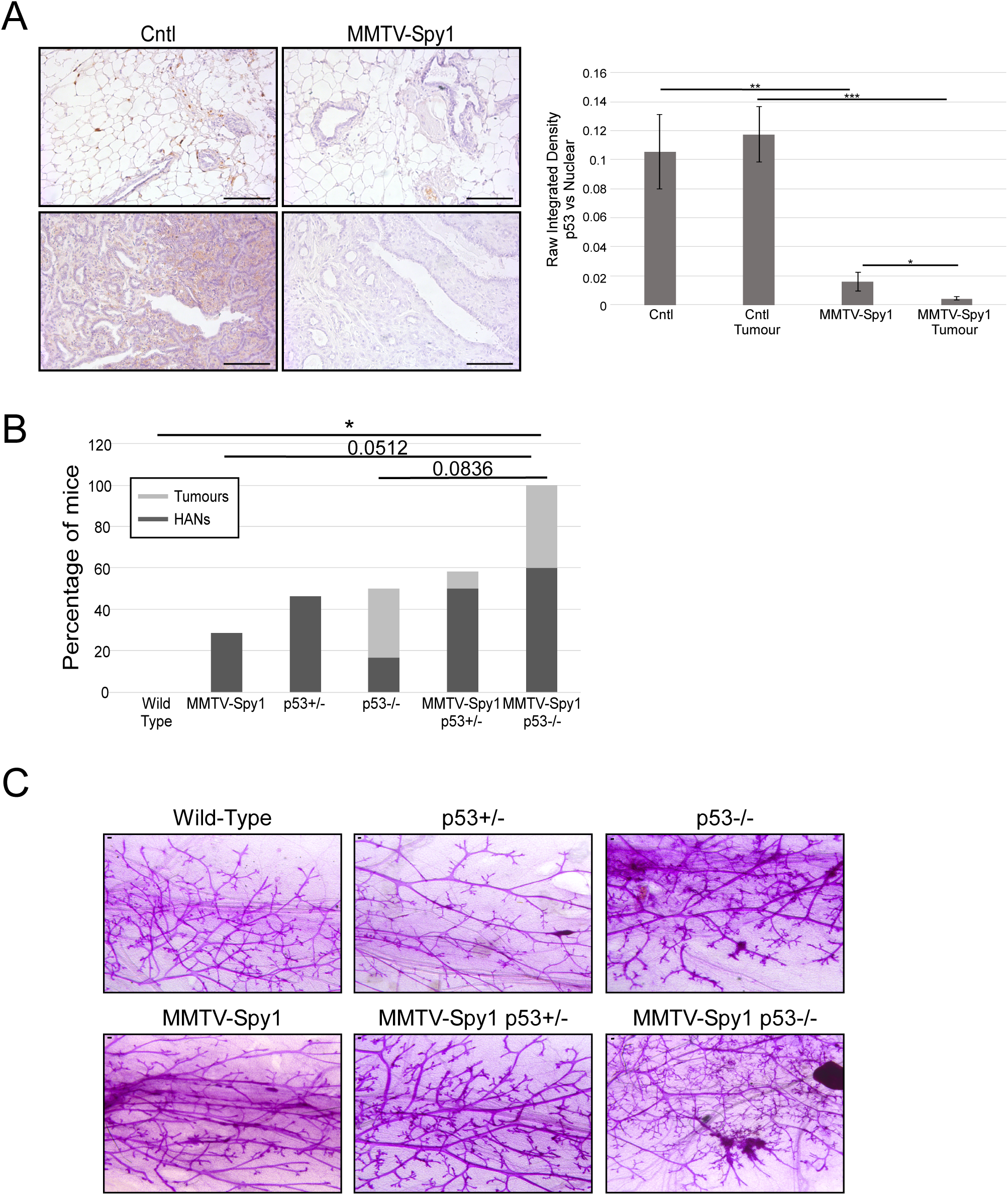
Loss of p53 enhances hyperplasia in MMTV-Spy1 mice. A) Immunohistochemical analysis for p53 expression in inguinal mammary glands and tumours of DMBA treated MMTV-Spy1 mice and littermate controls. Representative images are shown in left panel. Levels of p53 were quantified using ImageJ analysis (right panel). Scale bar=100 µm B) Fat pads of wild-type mice were reconstituted with mammary epithelial cells from MMTV-Spy1 mice crossed with p53 null mice and were monitored for HANs and formation of tumours. Only tumour negative mice were screen for the formation of HANs. (Wild-type n=5; MMTV-Spy1 n=7, p53+/- n=13; p53-/- n=6; MMTV-Spy1 p53+/- n=12; MMTV-Spy1 p53-/- n=5) C) Representative images of whole mounts. Scale bar=0.1mm. Errors bars represent SE; Student’s T-test (A), Mann-Whitney (B). *p<0.05, **p<0.01, ***p<0.001.

## Discussion

Development of the transgenic MMTV-Spy1 mouse has yielded new insight into the molecular regulation of the breast during development, revealing how misregulation of cell cycle checkpoints can impact susceptibility to tumorigenesis. On the tumor resistant B6CBAF1/J background the MMTV-Spy1 mice develop normally, showing no overt phenotypic differences and no spontaneous tumorigenesis, despite a significant increase in proliferative potential of mammary epithelial cells [40] Primary mammary epithelial cells also demonstrate increased proliferative potential. Previous data demonstrated that overexpression of Spy1 in the murine HC11 cell line shows disrupted two-dimensional acinar development *in vitro*, accelerated ductal development *in vivo*, and increased tumourigenesis when transplanted into cleared mammary fat pads [20]. One difference between these systems is the HC11 cell line contains a mutated p53 which renders p53 non-functional [36–38]. Investigating this hypothesis, we found that that knockdown of p53 in MMTV-Spy1 primary mammary epithelial cells increases Spy1 protein levels significantly. To examine the relationship between Spy1 and p53, we turned our attention to *in vitro* cell systems, using a variety of cell lines differing in the status of p53 and DNA repair pathways. We found an inverse relationship between Spy1 and p53 protein levels in every cell system studied, and constitutive induction of Spy1 was capable of abrogating p53 mediated effects on proliferation in all scenarios. This supports previous functional data demonstrating that Spy1 can override the DDR and bypass checkpoint responses [12, 13, 15, 16]. We also demonstrated that p53 mediated degradation of Spy1 is proteasome dependent and specifically requires the E3 ligase Nedd4. Collectively, these data support that p53 targets Spy1 protein levels to ensure the normal functioning of the DDR.

Mice treated with DMBA had elevated p53 levels, along with a significant increase in the number of γH2AX cells. The elevated p53 seen in the MMTV-Spy1 mice upon exposure to DMBA without the subsequent decrease in Spy1 levels shown in cell systems may be due to the strong viral promoter in the transgene which would allow for consistent elevation of Spy1 despite the mounting p53 response to try and decrease levels. Increased levels of γH2AX can signify latent unrepaired damage, or perhaps a delay in the repair response to DNA damage. Increased expression of γH2AX is indicative of increased levels of DNA damage, which in turn can lead to accumulation of deleterious mutations and onset of tumourigenesis. Alterations in the accumulation and subsequent decrease in γH2AX is also shown *in vitro* indicating alterations to the DNA damage response. We demonstrate that indeed the MMTV-Spy1 mice present with a significant increase in tumour formation. While there were some interesting findings with the histology of DMBA induced tumours, no significant differences were found between DMBA induced tumours in control versus MMTV-Spy1 mice. Many of the histologies noted are commonly found in DMBA induced tumours, however, further investigation is warranted to determine if Spy1 is capable of driving different subtypes or histologies of breast cancer [41, 42].

When crossed with p53 null mice, fat pads of wild-type mice reconstituted with mammary epithelial cells from intercrossed MMTV-Spy1 mice with loss of p53 had more hyperplasia and tumours over wild-type mice reconstituted with wild-type mammary epithelial cells. The data suggests that complete loss of p53 may enhance the ability of Spy1 to drive tumourigenesis. To test this MMTV-Spy1 primary mammary epithelial cells were manipulated for p53 levels and data supports this hypothesis, there is a significant increase in proliferation in the absence of p53. Future work to combine this with known oncogenic drivers is an important next step. Reports in the literature show the loss of p53 alone on a susceptible strain of mouse leads to formation of mammary tumours in 75% and 55% of p53 null and heterozygous mice respectively [43]. It is important to note the differences in strain between the reported literature and the MMTV-Spy1 and p53 intercross described in this study. While Balb/C mice are known to be more susceptible to mammary tumour formation, C57BL/6 mice are known to be more resistant, which may also account for lower rates of tumour onset seen with the MMTV-Spy1 and p53 null intercross [40, 44]. Our data supports that elevated protein levels of Spy1 cooperate with these events.

Increased susceptibility to breast cancer, such as familial cases of breast cancer, are often caused by inherited mutations in genes that regulate the DDR, such as BRCA and p53 [5, 11, 45, 46]. It is likely that other genes which mediate cell cycle progression and alter the DDR may also be involved in enhanced susceptibility. Interestingly, studies investigating genes involved in breast cancer susceptibility have identified chromosome 2p, and specifically 2p23.2, as a site which may have genes that contribute to increased breast cancer risk [47–49]. This identified location maps directly to the chromosomal location of the Spy1 gene (*SPDYA*). While further work is needed to definitively identify Spy1 as a breast cancer susceptibility gene, the current data provides support for Spy1 in enhancing susceptibility.

## Conclusions

Collectively, this work presents a novel feedback loop between the atypical cell cycle regulator Spy1 and the tumour suppressor protein p53, where tight control over Spy1 protein levels is required to maintain normal expansion of the developing mammary epithelium. When p53 is mutated, or Spy1 is expressed at elevated levels, this will allow for deleterious mutations to accumulate, increasing susceptibility to tumourigenesis (Figure 7). Restoring p53 function has been an elusive target in the clinic. Spy1-Cdk regulation is a unique and potentially potent mechanism for drug design, which may represent a novel therapeutic approach for select forms of breast cancer.

**Figure 7.**
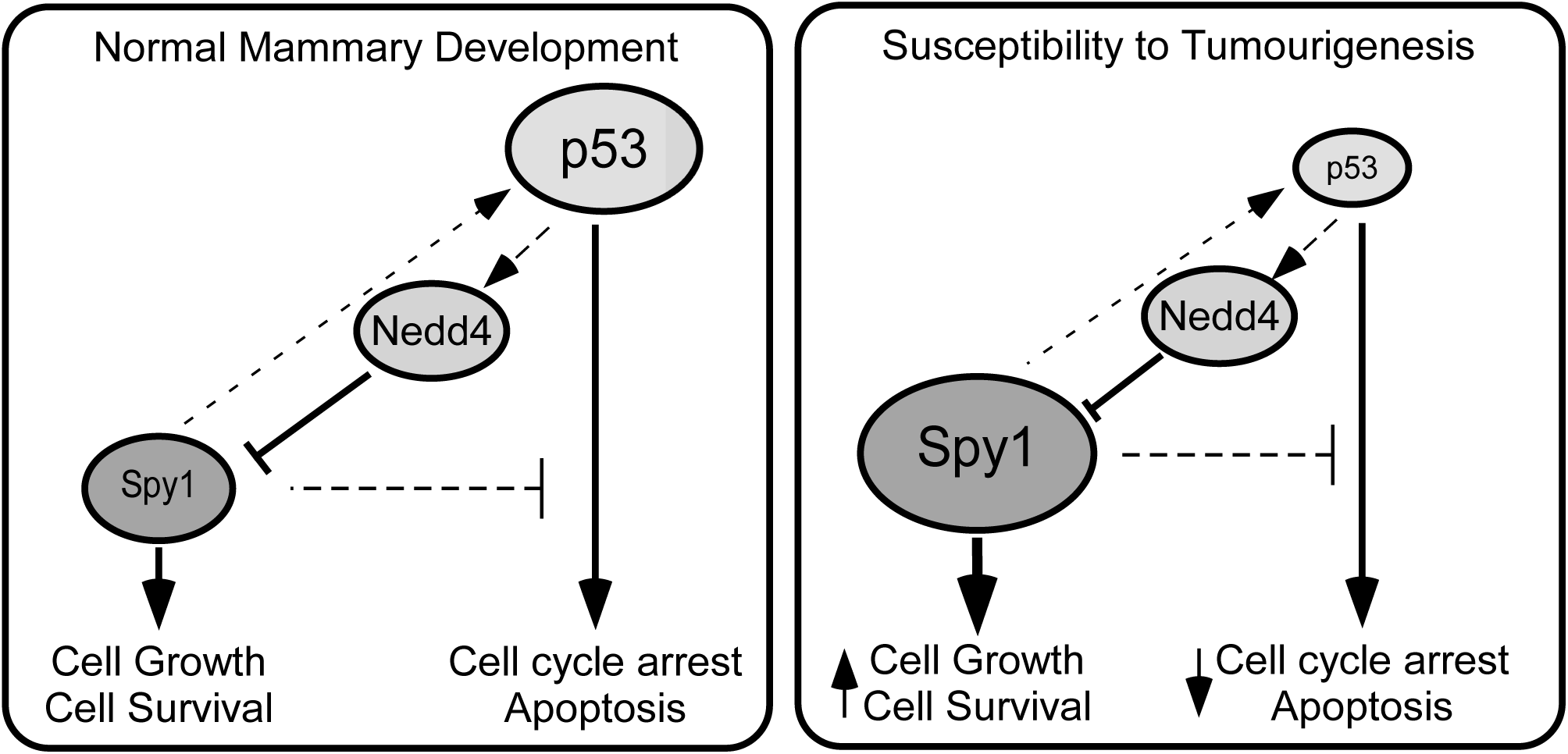
Mechanism for increased susceptibility by elevation of Spy1. Left panel reflects that Spy1 protein levels are held in check by wild-type p53 to allow tightly regulated bursts of needed mammary proliferation during development. The panel to the right reflects the situation when either p53 is mutated/deleted or Spy1 protein levels are elevated, supporting susceptibility to tumorigenesis.

## Ethics Approval and Consent to Participate

All experiments performed were approved by the University of Windsor Animal Care Committee.

## Consent for Publication

All authors have agreed to publish this manuscript.

## Availability of data and materials

All data generated from this study are included in the manuscript and additional file 1 supplemental files.

## Author Contributions

BF, IQ, EK and LAP contributed to project design. BF, IQ, EK and RDC contributed to data acquisition. BF, IQ, RDC and LAP contributed to data analysis. BF, IQ and LAP prepared the manuscript. LAP secured the funding for this study.

## Acknowledgements

We thank Drs. C. Pin, F. Dick and L. Drysdale from the London Regional Transgenic and Gene Targeting Facility for the transgenic injections and helpful advice. Drs. M. Crawford and D. Higgs for use of equipment. Special thanks to Dr. W. Muller for the MMTV-SV40-TRPS-1 vector and Dr. C. Shemanko for donation of HC11 cell line. Thanks to A. Malysa, E. Jalili, D. Lubanska, E. Laurie, M. Elliot and N. Paquette for assistance with statistics, vector construction, idea generation, genotyping and technical assistance.

## Supplemental Files

Additional file 1: Figure S1.

Additional file 2: Figure S2.

Additional file 3: Figure S3.

Additional file 4: Figure S4.

Additional file 5: Figure S5.

Additional file 6: Supplemental Figure Legends

