## Supplementary figures and images for "The Atypical Cyclin-Like Protein Spy1 Overrides p53-Mediated Tumour Suppression and Promotes Susceptibility to Breast Tumorigenesis"

### Supplemental Figures

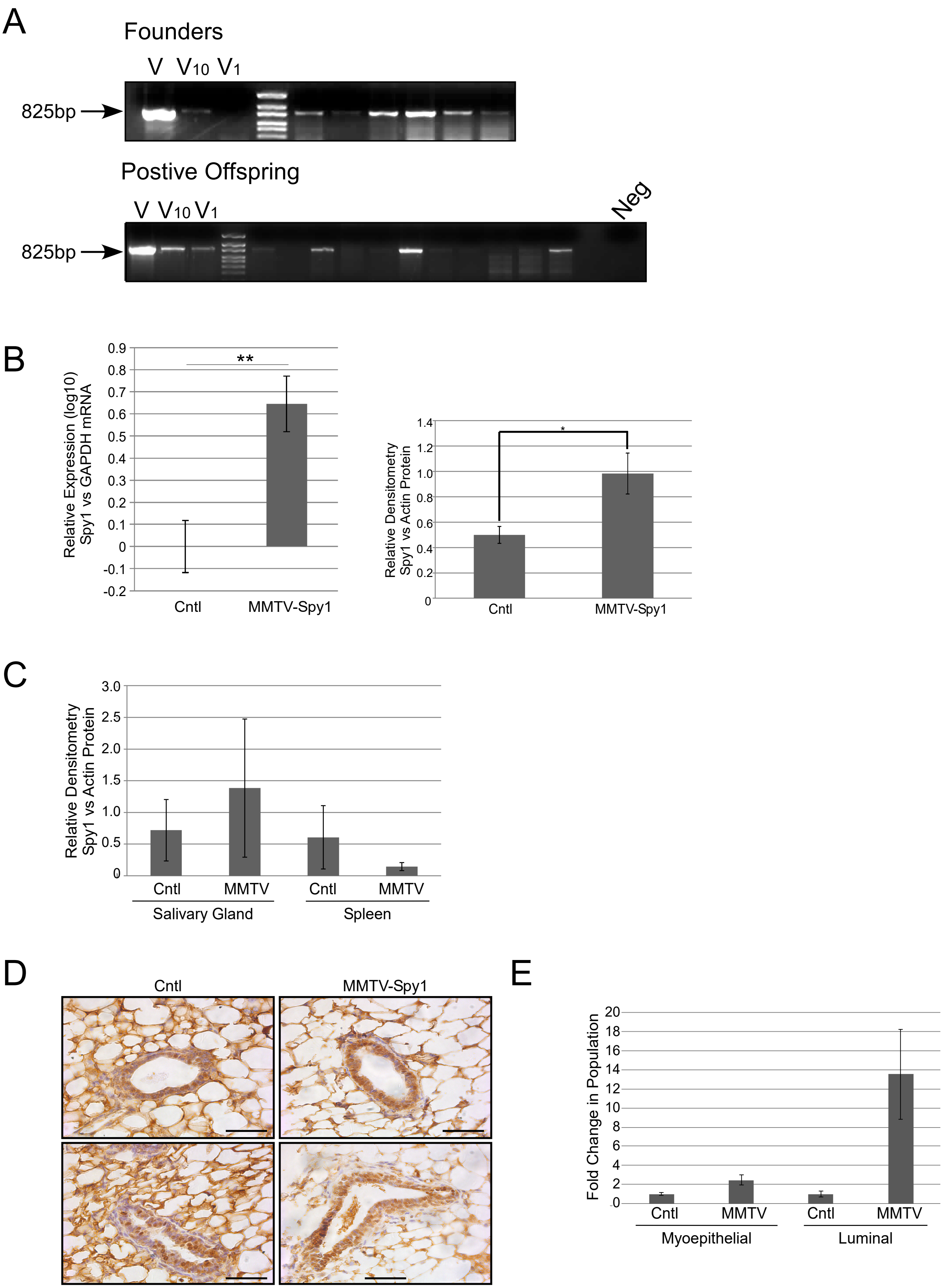

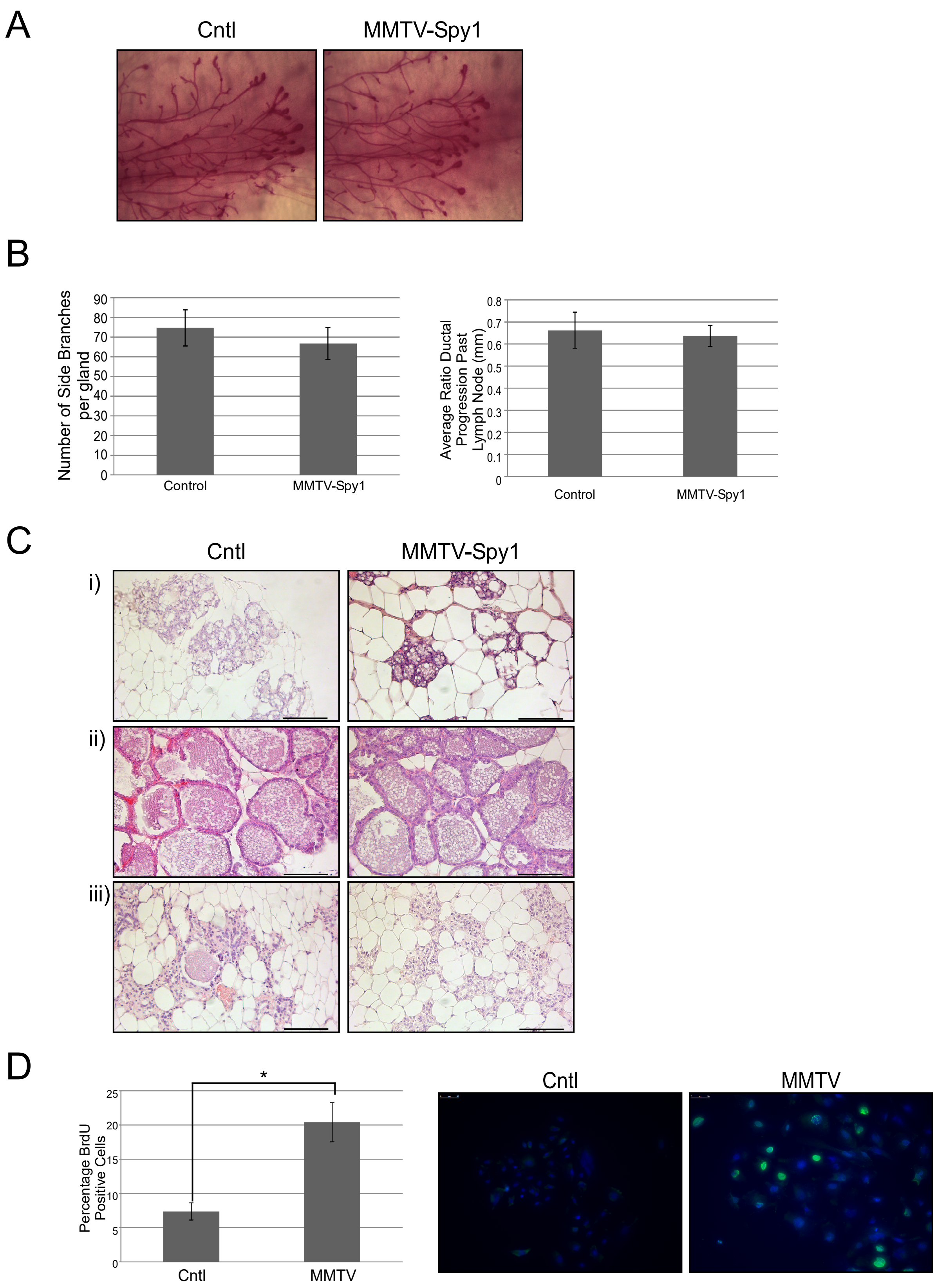

Figure S3

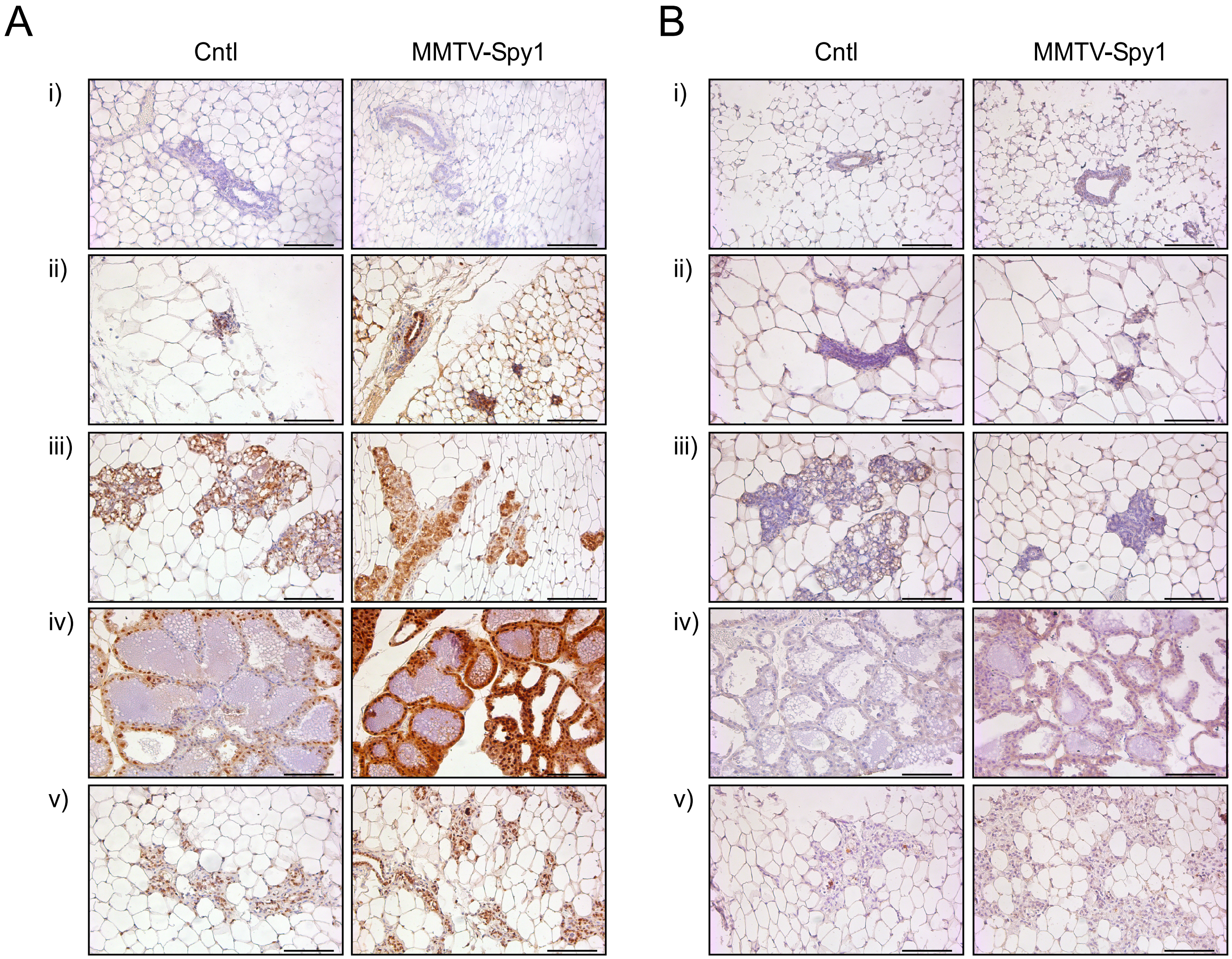

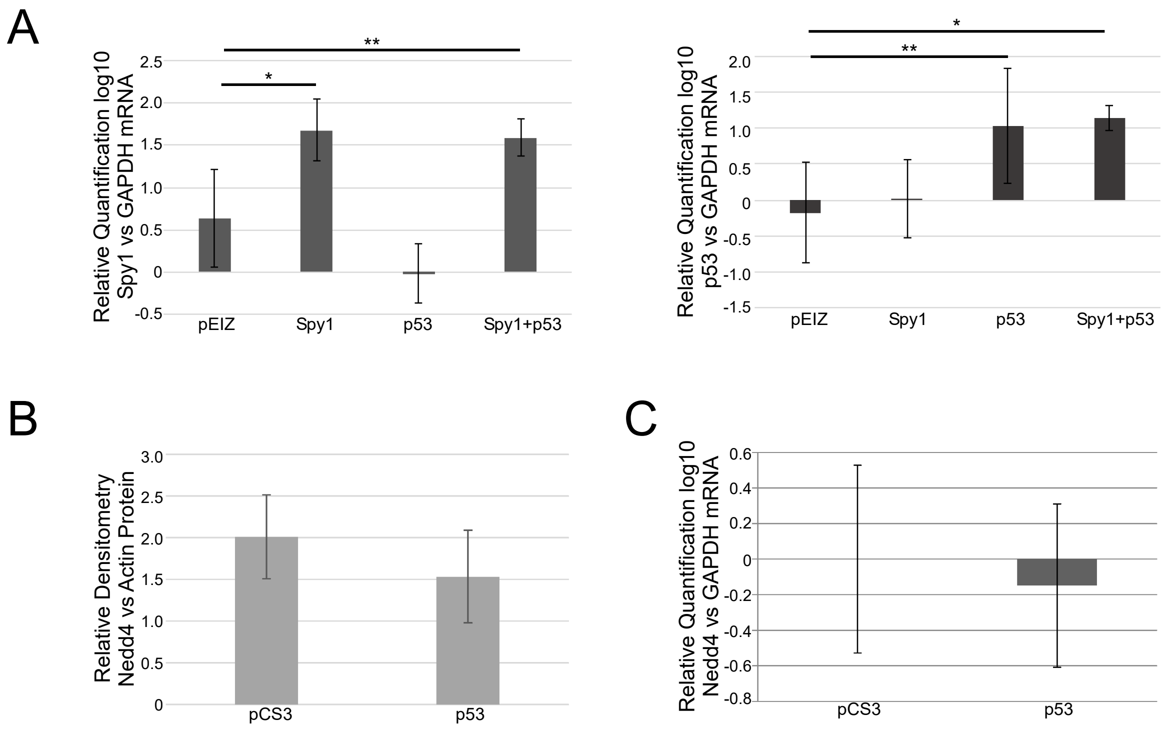

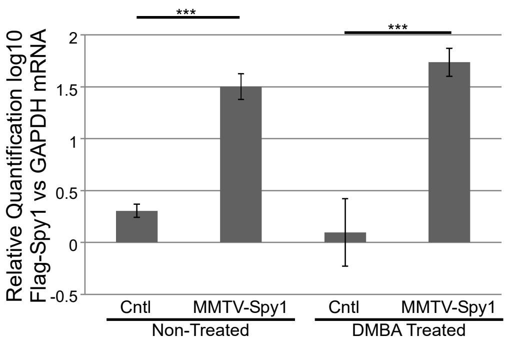

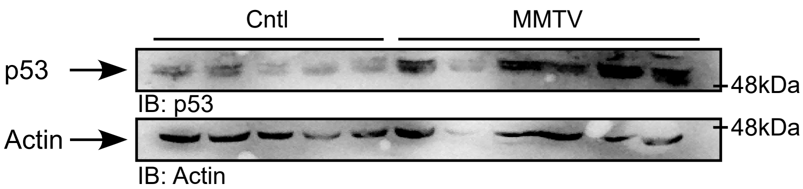
