## Supplemental Figure Legends for "The Atypical Cyclin-Like Protein Spy1 Overrides p53-Mediated Tumour Suppression and Promotes Susceptibility to Breast Tumorigenesis"

**Supplementary Figures**

**Figure S1; Related to Figure 1:** Characterization of MMTV-Spy1 mouse model system. A) Blots showing PCR analysis to confirm presence of transgene in founder mice (upper blot) and in offspring from founders (lower blot) where V represents vector control, V10 represents 10 copy vector control and V1 represents 1 copy vector control. B) qRT-PCR analysis of MMTV-Spy1 and littermate control (cntl) inguinal gland samples for Spy1 levels corrected for total levels of GAPDH. (n=8; left panel). Levels of Spy1 protein in 6-week-old MMTV-Spy1 inguinal glands were quantified and corrected for total actin protein levels (right panel) (n=8). C) Quantification of densitometry analysis of Spy1 protein levels corrected for total actin levels (right panel). (Salivary gland n=3; Spleen n=6). D) Representative images of Spy1 immunohistochemical analysis in MMTV-Spy1 and littermate control 8-week-old mice showing Spy1 localization within the mammary gland. Scale bar=50 µm. E) Flow cytometry of primary cells extracted from MMTV-Spy1 and littermate controls and stained for CD24 and CD45. Fold change in myoepithelial ($CD24^{lo}CD45^{-}$) and luminal ($CD24^{hi}CD45^{-}$) cell population is depicted (n=3). Error bars reflect SE. Student's T-test *p<0.05, **p<0.01

**Figure S2; Related to Figure 1:** Analysis of MMTV-Spy1 early development. A) Representative images of whole mount analysis from 6-week-old MMTV-Spy1 and littermate control (cntl) B6CBAF1/J mice (Cntl n=4, MMTV-Spy1 n=4). B) Graphical representation of analysis of whole mount images from B6CBAF1/J inguinal glands from MMTV-Spy1 and littermate controls. Number of side branches per gland was quantified and the average number of side branches per gland was calculated (left panel). Ratio of ductal progression of ductal network past the lymph node was measured for each gland,

and the average rate of ductal progression is shown in the right panel. C) Representative hematoxylin and eosin images from i) 16.5 day pregnancy, ii) 4 day lactation and iii) 4 day involution. Scale bar= 100 µm. D) Representative images (right panel) and quantification of BrdU incorporation (left panel) of primary mammary epithelial cells isolated from MMTV-Spy1 mice and their control littermates (cntl) (n=4 separate isolations). Scale bar= 50 µm. Error bars represent SE. *$p<0.05$

**Figure S3; Related to Figure 1:** Spy1 increases proliferation during development in mammary epithelial cells. Representative images are shown of A) PCNA and B) cleaved caspase 3 at i) 8 week puberty, ii) 12 week adult, iii) 16.5 day pregnancy, iv) 4 day lactation and v) 4 day involution. Scale bar= 100 µm.

**Figure S4. Related to Figure 3.** A) Spy1 and p53 were overexpressed in MDA-MB-231 cells to determine if p53 can alter Spy1 mRNA levels (n=3). B & C) p53 or control vector was overexpressed in HEK-293 cells to assess protein and RNA levels of Nedd4. B) Western blot analysis of Nedd4 protein levels corrected for total Actin protein levels. C) qRT-PCR analysis of Nedd4 RNA levels corrected for total GAPDH. Error bars represent SE. *$p<0.05$, **$p<0.01$

**Figure S5. Related to Figure 5.** A) qRT-PCR analysis of Spy1 levels in 8-week-old MMTV-Spy1 mice and their control littermates (cntl) 48 hours after DMBA treatment in mice with and without DMBA. Levels of Flag-Spy1 are corrected for total levels of GAPDH. B) Representative western blot for p53 protein levels in MMTV-Spy1 8-week-old mice and their control littermates 48 hours after DMBA treatment. Error bars represent SE. ***$p<0.001$
